# Reprogramming of Iron Metabolism Confers Ferroptosis Resistance in ECM-Detached Cells

**DOI:** 10.1101/2022.09.23.509253

**Authors:** Jianping He, Abigail M. Abikoye, Brett P. McLaughlin, Ryan S. Middleton, Ryan Sheldon, Russell G. Jones, Zachary T. Schafer

## Abstract

Cancer cells often acquire resistance to cell death programs induced by loss of integrin-mediated attachment to extracellular matrix (ECM). Given that adaptation to ECM-detached conditions can facilitate tumor progression and metastasis, there is significant interest in effective elimination of ECM-detached cancer cells. Here, we find that ECM-detached cells are remarkably resistant to the induction of ferroptosis. While alterations in membrane lipid content are observed during ECM-detachment, it is instead fundamental changes in iron metabolism that underlie resistance of ECM-detached cells to ferroptosis. More specifically, our data demonstrate that levels of free iron are low during ECM-detachment due to changes in both iron uptake and iron storage. In addition, we establish that lowering the levels of iron storage proteins sensitizes ECM-detached cells to death by ferroptosis. Taken together, our data suggest that therapeutics designed to kill cancer cells by ferroptosis may be hindered by lack of efficacy towards ECM-detached cells.

## Introduction

Mammalian cells often maintain integrin-mediated attachment to the extracellular matrix (ECM) in order to antagonize cell death programs (Buchheit et al., 2012). More specifically, loss of ECM-attachment is well-known to induce a cell death program termed anoikis, which involves caspase-dependent cell death by apoptosis (Frisch and Francis, 1994). However, ECM-detachment is also well recognized to induce numerous, additional changes that can impact cell viability independent from anoikis (Mason et al., 2017). These changes include profound alterations in nutrient uptake (Grassian et al., 2011; Schafer et al., 2009), metabolic reprogramming (Jiang et al., 2016; Mason et al., 2021; Mason et al., 2016), the induction of mitophagy (Hawk et al., 2018), and the rewiring of signal transduction (Buchheit et al., 2015; Debnath et al., 2002; Overholtzer et al., 2007; Reginato et al., 2005; Reginato et al., 2003; Tsegaye et al., 2021; Weigel et al., 2014). Given that cancer cells often encounter ECM-detached conditions during tumor progression and metastasis (Buchheit et al., 2014), the mechanisms by which cancer cells evade ECM-detachment-mediated cell death programs could ultimately serve as targets for the elimination of invasive cancer cells.

While we (and others) have long been interested in understanding how cancer cells adapt and survive during ECM-detachment, less attention has been devoted to attempts to specifically kill cancer cells that have successfully adapted to ECM-detached conditions. In addition to the therapeutic possibilities that could be unveiled by such investigations, understanding the sensitivity (or insensitivity) of ECM-detached cells to extrinsic treatments could also generate important new biological insight. One such avenue that has been explored as a strategy to eliminate cancer cells is the induction of ferroptosis, an iron-dependent, non-apoptotic form of programmed cell death that is characterized by robust lipid peroxidation (Dixon et al., 2012; Jiang et al., 2021; Magtanong et al., 2022; Stockwell, 2022). Sensitivity to cell death by ferroptosis can be regulated by diminished glutathione synthesis owing to impaired cystine uptake (Badgley et al., 2020; Kremer et al., 2021; Poursaitidis et al., 2017; Zhang et al., 2021), alterations in the abundance or activity of the antioxidant selenoprotein glutathione peroxidase 4 (GPX4) (Yang et al., 2014; Yang and Stockwell, 2008), or alterations in membrane lipid content that favor oxidation (Doll et al., 2017; Wu et al., 2019). In addition to these regulatory pathways, recent studies have continued to unveil additional factors that impact the activation or inhibition of ferroptosis (Bersuker et al., 2019; Brown et al., 2019; Doll et al., 2019; Li et al., 2022; Mishima et al., 2022; Venkatesh et al., 2020). Currently, there is significant interest in the possibility that induction of ferroptosis may be an attractive strategy to kill cancer cells and thus limit tumor progression (Gao et al., 2021; Hassannia et al., 2019; Koppula et al., 2022; Tarangelo et al., 2018; Wiernicki et al., 2022; Wu et al., 2022; Yi et al., 2020; Zhang et al., 2018). However, the sensitivity of ECM-detached cells to ferroptosis induction has not been investigated in detail.

Here, we demonstrate that ECM-detached cells are strikingly resistant to the induction of ferroptosis. This protection from ferroptosis induction extends across multiple cell lines and to both pharmacological and genetic means of ferroptosis activation. ECM-detached cells are well-known to form multicellular aggregates and enhanced cell-cell contacts are linked to alterations in lipid content that cause ferroptosis resistance. However, aggregation-induced changes in membrane lipid content do not underlie ferroptosis resistance during ECM-detachment. Instead, we find that ECM-detached cells have low levels of free (redox-active) iron and that addition of excess iron is sufficient to sensitize ECM-detached cells to ferroptosis induction. We also find that iron uptake and iron storage are altered during ECM-detachment and that limiting the levels of iron storage proteins can sensitize ECM-detached cells to ferroptosis. Taken together, our results suggest a direct link between ECM-detachment and iron metabolism. Strategies that attempt to kill cancer cells through ferroptosis may be hindered as a consequence of this intrinsic resistance during ECM-detachment.

## Results

### ECM-detached cells are resistant to pharmacological and genetic induction of ferroptosis

To examine the sensitivity of ECM-detached cells to the induction of ferroptosis, we treated various cell lines with increasing concentrations of erastin, which triggers ferroptosis through the inhibition of the system x_c_^-^ antiporter (Dixon et al., 2012). We intentionally selected cell lines derived from disparate tissue types to assess whether any observed changes in erastin sensitivity were fundamentally distinct in specific cellular contexts. Interestingly, we found that when 786-O, RCC10, MDA-MB-231, SUM159, and HeLa cells were grown in ECM-detached conditions, they were strikingly resistant to erastin-induced cell death (Figs. 1A, S1A). The effect of system x_c_^-^ antiporter inhibition (and consequent ferroptosis induction) can also be achieved by elimination of cystine from the cell culture media (Badgley et al., 2020; Kremer et al., 2021; Poursaitidis et al., 2017; Zhang et al., 2021). Indeed, ECM-detached cells, unlike their ECM-attached counterparts, are resistant to cystine-starvation induced cell death (Fig. S1C). We also tested the sensitivity of ECM-detached cells to RSL3, which is known to induce ferroptotic cell death by blocking the activity of GPX4 (Yang et al., 2014; Yang and Stockwell, 2008). Indeed, the capacity of RSL3 to induce cell death is markedly compromised when these cell lines are grown in ECM-detachment (Figs. 1B, S1B). As expected, treatment with ferrostatin-1 (Fer-1), a well-characterized inhibitor of ferroptosis (Dixon et al., 2012), led to a substantial inhibition of erastin-induced (Fig. 1C), RSL3-induced (Fig. 1D), and cystine starvation-induced (Fig. S1C) cell death in ECM-attached cells. However, the viability of ECM-detached cells treated with these stimuli was not meaningfully altered by Fer-1 treatment. To assess whether the resistance of ECM-detached cells to ferroptosis is reversible, we grew cells in ECM-detached conditions before allowing them to re-attach to the plate. Indeed, the resistance to erastin-induced death acquired during ECM-detachment was lost upon reattachment to extracellular matrix (Fig. S1D).

**Figure 1.**
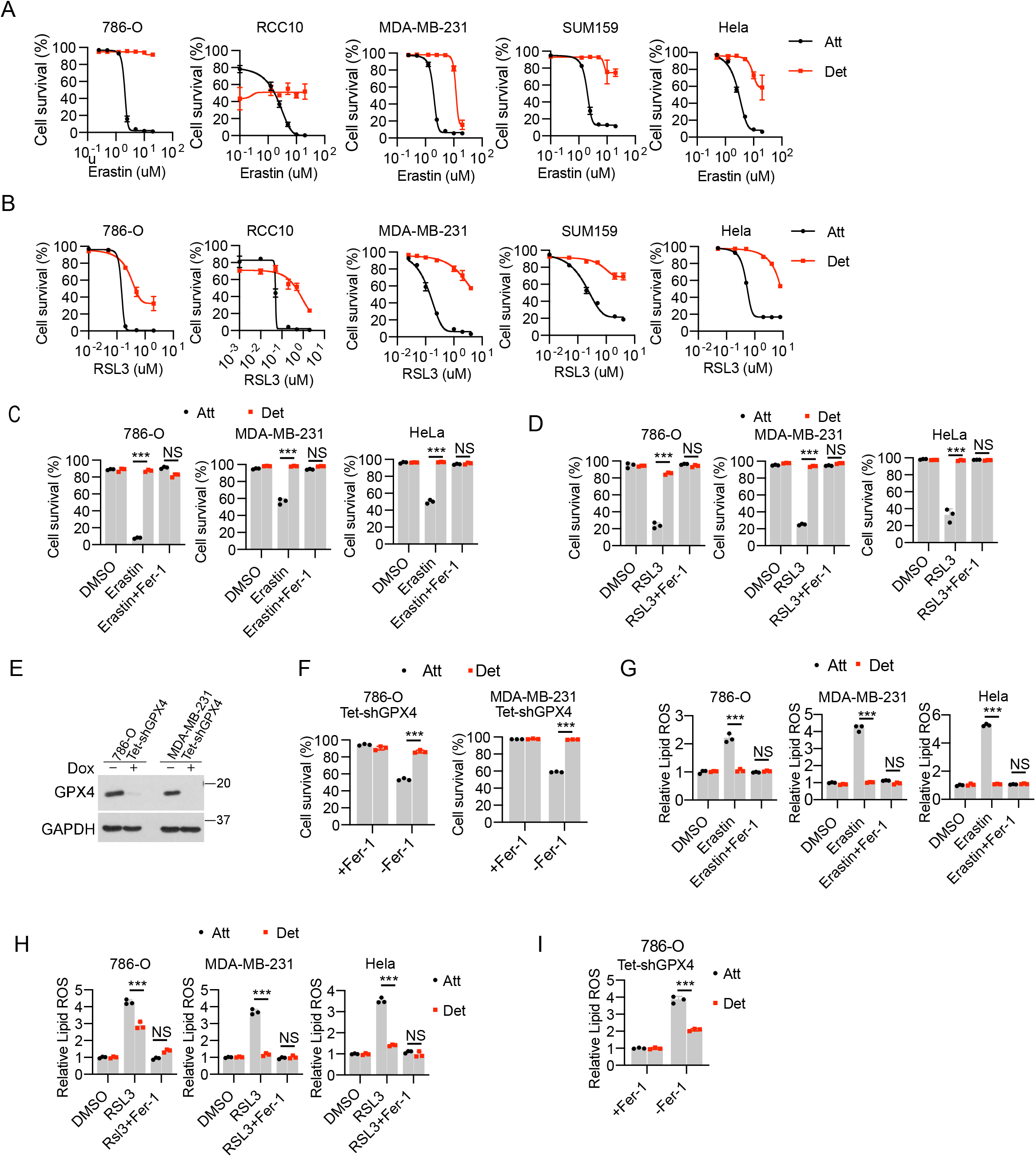
ECM-detached cells are resistant to ferroptosis induction. **(A-B)** Survival of various tumor cell lines with erastin treatment for 24 h (A) or RSL3 treatment for 18 h (B). **(C-D)** Survival of 786-O, MDA-MB-231 and Hela cells to erastin (5 μM, 24 h) or RSL3 (0.2 μM, 12 h). Ferrostatin-1 (Fer-1) was used at 2.5 μM. **(E)** Western blot to verify doxycycline inducible GPX4 knockdown in 786-O and MDA-MB-231 cells. Cells were kept in medium containing 2.5 μM Fer-1. **(F)** Cell survival after 20-h withdrawal of Fer-1. **(G-I)** Lipid ROS detection by C11 BODIPY 581/591 staining in erastin (5 μM, 12 h) (G), RSL3 (0.2 μM, 3 h) (H) or Fer-1 withdrawal (5 h) (I) treatments. Fer-1 (2.5 μM) was used to block ferroptosis. ***p < 0.001; NS, not significant; two-way ANOVA followed by Tukey test. Data are mean ±SD. Graphs represent data collected from a minimum of three biological replicates and all western blotting experiments were independentlyrepeated a minimum of three times with similar results.

To complement these studies using a genetic method of ferroptosis induction, we utilized cells engineered to express doxycycline-inducible GPX4 shRNA (Fig. 1E). These cells grow in the presence of Fer-1 and undergo ferroptosis upon removal of Fer-1 from the media (Friedmann Angeli et al., 2014). Unlike when cells are grown in ECM-attached conditions, Fer-1 withdrawal in cells deficient in GPX4 failed to induce cell death in ECM-detached 786-O or MDA-MB-231 cells (Fig. 1F). Furthermore, the accumulation of lipid ROS as measured by C11 BODIPY 581/591 in response to erastin (Fig. 1G), RSL3 (Fig. 1H), cystine starvation (Figure S1E), or GPX4 shRNA (Fig. 1I) treatment is blunted when cells are grown in ECM-detachment. Additionally, lipid ROS accumulation in ECM-detached cells exposed to erastin was restored upon reattachment of ECM-detached cells to the plate (Fig. S1F). Importantly, the resistance attained in ECM-detached cells does not extend to more generic oxidative stress as we do not observe differences between ECM-detached and –attached cells in the sensitivity of H_2_O_2_ treatment (Fig. S1G). In aggregate, these results suggest that cells grown in ECM-detached conditions are resistant to ferroptosis induction by pharmacological and genetic means.

### Loss of Yap-mediated ACSL4 does not underlie ferroptosis resistance during ECM-detachment

We next sought to assess the molecular mechanism by which ECM-detached cells are resistant to the induction of ferroptosis. Previous studies in cancer cells suggest that growth in ECM-detached conditions (including as circulating tumor cells) can result in cell-cell clusters that promote cell survival and ultimately, tumor progression and metastasis (Aceto et al., 2014; Brown et al., 2018; Cheung et al., 2016; Liu et al., 2019; Rayavarapu et al., 2015). Similarly, the formation of cell-cell clusters has also been linked to ferroptosis resistance due to the inhibition of Yap-mediated expression of ACSL4 (Wu et al., 2019), a protein that stimulates ferroptosis by promoting the incorporation of oxidizable polyunsaturated fatty acid phospholipids (PUFA-PLs) into membranes (Doll et al., 2017). Given these findings, we sought to assess whether aggregation-induced changes in lipid composition were underlying the resistance of ECM-detached cells to ferroptosis. We first conducted mass spectrometry-based lipidomics analysis and observed alterations in a number of distinct lipid species as a consequence of growth in ECM-detachment (Fig. 2A). More specifically, growth in ECM-detached conditions resulted in a significant loss of PUFA-containing phosphatidylcholine (PC) and phosphatidylethanolamine (PE) (Fig. 2B). In addition, levels of ACSL4 protein (Fig. 2C) and transcript (Fig. 2D) were substantially reduced in ECM-detached conditions. Notably, we did not observe large changes in the abundance of other ferroptosis regulators (GPX4, FSP-1, system x_c_^-^, GCLC) that were likely to account for the resistance of ECM-detached cells to ferroptosis induction (Fig. 2C). In sum, these results are consistent with the hypothesis that diminished abundance of ACSL4-mediated PUFAs may contribute to ferroptosis resistance during ECM-detachment.

**Figure 2.**
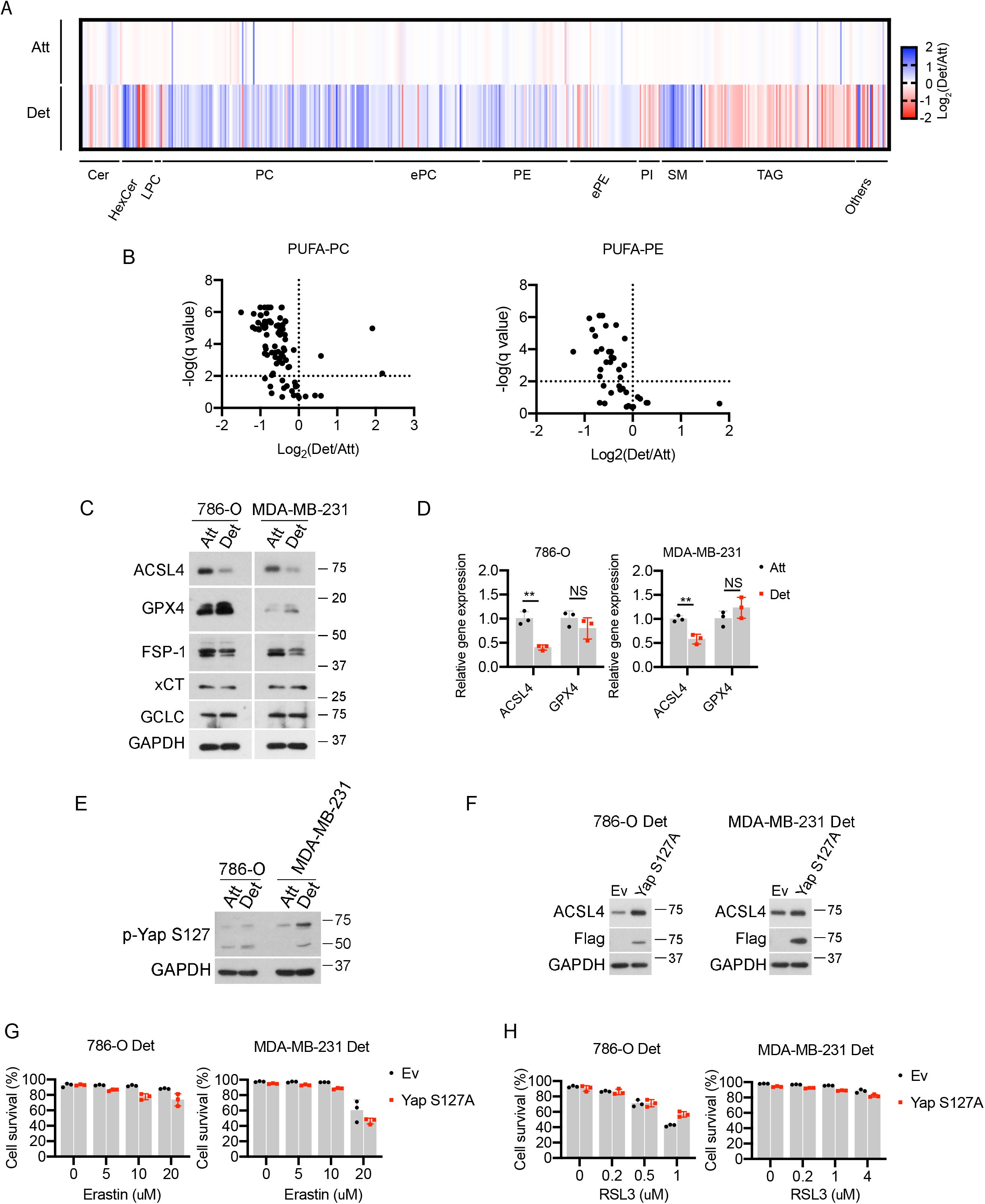
Loss of Yap-regulated ACSL4 does not account for ferroptosis resistance in ECM-detached cells. **(A)** Heatmap of the relative lipid abundance between attached and detached 786-O cells after 24-h detachment. Abbreviations: Cer, ceramide; HexCer: hexosylceramide; LPC, lysophosphatidylcholine; PC, phosphadylcholine; TAG, triacylglycerol; PE, phosphatidylethanolamine; ePE, (vinyl ether-linked) PE-plasmalogen; PI, phosphatidylinositol; SM, sphingomyelin. Blue: down-regulated relative to the attached cells, red: upregulated relative to the attached cells. **(B)** Volcano plot for poly-unsaturated fatty acid linked phosphatidylcholine (PUFA-PC) or phosphatidylethanolamine (PUFA-PE). **(C)** Western blot detection of protein levels in cells after 24-h detachment. **(D)** Gene expression by quantitative real-time PCR. **(E)** Western blot detection of p-Yap S127 after 24-h detachment. **(F)** Western blot to verify Yap S127A overexpression and rescue of ACSL4 expression under ECM-detachment. **(G-H)** Cell survival in empty vector (Ev) or Yap S127A expressing cells treated with increasing doses of erastin (G) or RSL3 for 24 h (H). **p < 0.001; NS, not significant; two-tailed student’s test. Data are mean ±SD. Graphs represent data collected from a minimum of three biological replicates and all western blotting experiments were independently repeated a minimum of three times with similar results.

As mentioned above, loss of ACSL4 has been previously linked to the diminished Yap activity as a consequence of cell-cell contacts. Indeed, consistent with previous findings (Wu et al., 2019), phosphorylation of Yap at serine 127 (which prevents Yap activation by blocking its translocation to the nucleus) was elevated when cells are grown in ECM-detachment (Fig. 2E). Yap activity can be enhanced through the generation of the S127A mutation which prevents phosphorylation and thus enhances nuclear retention (Wu et al., 2019). Indeed, expression of the Yap S127A mutant in ECM-detached cells results in elevated levels of ACSL4 (Fig. 2F). Despite the restoration of ACSL4 levels in ECM-detached cells, cells expressing Yap S127A remain similarly resistant to ferroptosis induction by erastin (Fig. 2G) or RSL3 (Fig. 2H). To complement these studies, we also disrupted cell-cell contacts directly using EDTA (Fig. S2A) or methylcellulose (MC) (Fig. S2B) and assessed the sensitivity to ferroptosis induction. While we did observe marginal changes in erastin or RSL3-mediated cell survival as a consequence of EDTA or MC treatment (Figs. S2A, S2B), the levels of lipid ROS remained low during ECM-detachment (Figs. S2C, S2D). Taken together these results suggest that alterations in the cell contact-Yap-ACSL4 axis do not underlie the sensitivity of ECM-detached cells to ferroptosis induction.

### ECM-detachment causes alterations in iron uptake

Given that the ECM-detachment-mediated changes in ACSL4 do not confer ferroptosis resistance and that we did not detect changes in the abundance of other proteins known to be involved in the detoxification of lipid ROS (Fig. 2C), we next investigated whether ECM-detachment resulted in modifications in iron metabolism. Interestingly, the levels of free (redox-active) iron were diminished during ECM-detachment when cells were grown in either normal or iron overload (FeCl3 addition) conditions (Fig. 3A). Similarly, free iron levels were markedly reduced when ECM-detached cells were treated with erastin (Fig. S3A) or RSL_3_ (Fig. S3B). Thus, we hypothesized that the lower levels of free iron in ECM-detached cells may contribute to the observed ferroptosis resistance. Indeed, supplementation with excess iron (FeCl_3_ addition) was sufficient to sensitize ECM-detached cells to death caused by RSL3 (Fig. 3B), erastin (Fig. 3C), or cystine starvation (Fig. S3C). Analogous results were obtained using a genetic method of ferroptosis induction as shRNA targeting GPX4 can cause cell death in ECM-detached cells when iron supplementation was provided (Fig. 3D).

**Figure 3.**
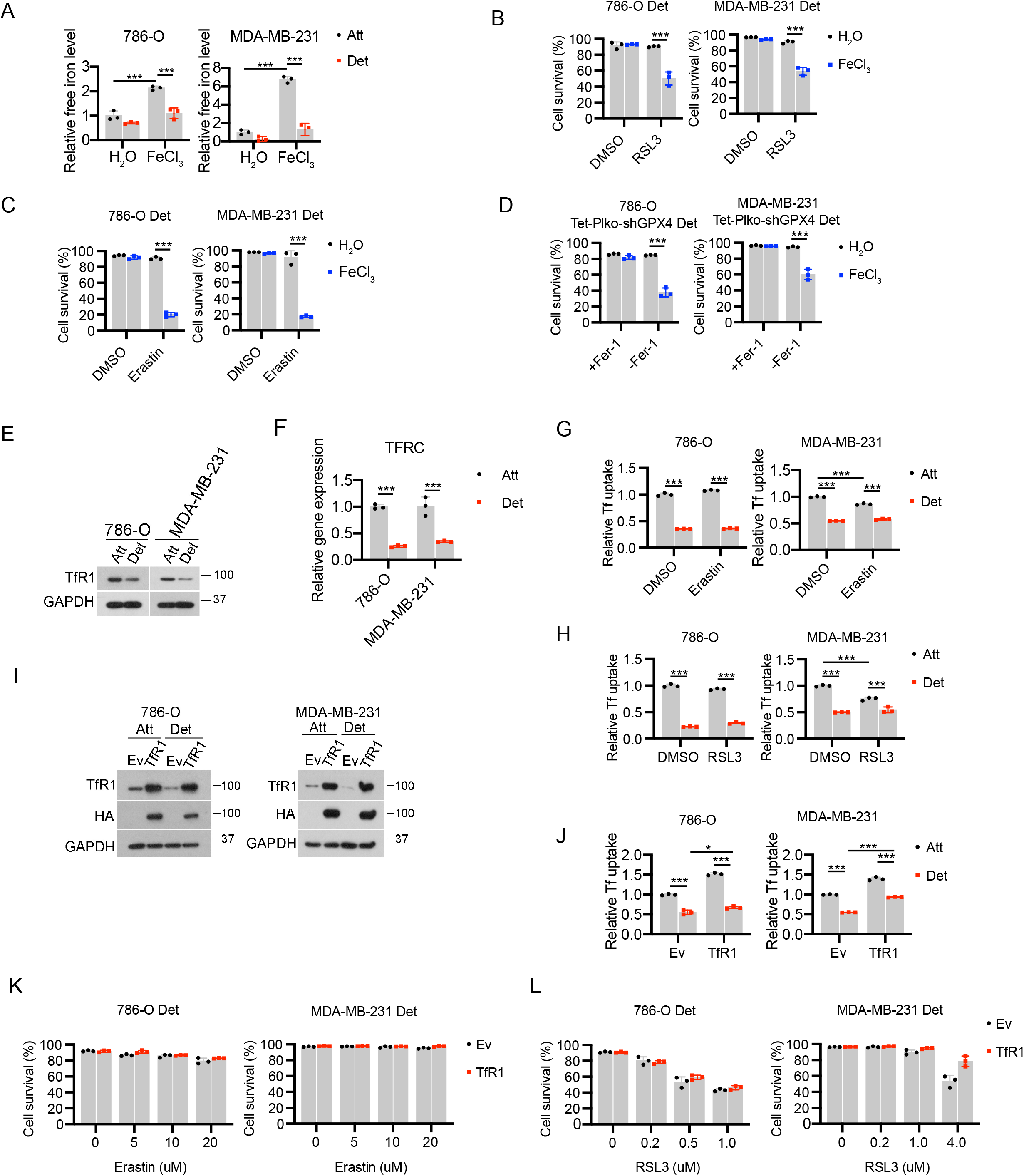
Iron uptake is reduced in ECM-detached cells. **(A)** Free iron level detection in cells treated with either H2O or FeCl3 (100 μM) for 24 h. **(B-D)** Cell survival in detached cells treated with erastin (B), RSL3 (C) or Fer-1 withdrawal (D) together with either H2O or FeCl3 (100 μM) for 24 h. **(E-F)** TfR1 protein (E) and mRNA (F) expression in cells detached for 24 h. **(G-H)** Transferrin uptake assay in 786-O and MDA-MB-231 cells treated with erastin (5 μM, 12 h) (G) or RSL3 (0.2 μM, 3 h) (H). **(I)** Western blot detection of TfR1 in TfR1-overexpressing cells. **(J)** Transferrin uptake assay in Ev or TfR1 overexpressed cells. **(K-L)** Cell survival in detached cells treated with increasing doses of erastin (K) or RSL3 (L) for 24 h. ***p < 0.001, two-tailed student’s t test in F. *p < 0.05, ***p < 0.001, two-way ANOVA followed by Tukey test in A, B, C, D, G, H, and J. Data are mean ±SD. Graphs represent data collected from a minimum of three biological replicates and all western blotting experiments were independently repeated a minimum of three times with similar results.

We next sought to determine why ECM-detached cells have lower levels of free-iron than their ECM-attached counterparts. As such, we reasoned that alterations in iron uptake may account for the observed differences. Indeed, we discovered that ECM-detachment resulted in lower levels of transferrin receptor 1 protein (TfR1, which internalizes transferrin-bound iron through clathrin-mediated endocytosis (Shen et al., 2018)) as measured by western blot (Fig. 3E) or qRT-PCR (Fig. 3F). We also found that transferrin uptake was reduced in ECM-detached conditions in the presence and absence of erastin (Fig. 3G) or RSL3 (Fig. 3H). These data suggest that deficiencies in transferrin-mediated iron uptake could contribute to the resistance of ECM-detached cells to ferroptosis induction. To test this possibility, we engineered 786-O or MDA-MB-231 cells to express high levels of HA-tagged TfR1 and measured the sensitivity of these cells to ferroptosis induction during ECM-detachment. Despite robust elevation of TfR1 in our cell lines (Fig. 3I) and a concurrent increase in transferrin uptake (Fig. 3J), we did not observe appreciable differences in erastin (Fig. 3K) or RSL3 (Fig. 3L)-mediated death in cells overexpressing TfR1. These results suggest that rescuing TfR1 levels is not sufficient to sensitize ECM-detached cells to ferroptosis-inducing agents.

### Alterations in FTH1 contribute to ferroptosis resistance during ECM-detachment

Given that rescue of TfR1 levels is not sufficient to sensitize ECM-detached cells to ferroptosis induction, we sought to investigate whether changes in iron storage were altered as a consequence of growth in ECM-detachment. Most cells will store excess iron in ferritin, a protein made up of ferritin heavy chain (FTH1) and ferritin light chain (FTL) subunits (Torti and Torti, 2013). Indeed, the levels of both FTH1 and FTL protein (Fig. 4A) and transcript (Fig. 4B) were increased upon growth in ECM-detachment. The elevated levels of FTH1 during ECM-detachment were maintained in iron replete (FeCl_3_ treated) but blocked during iron deficient (DFO treated) conditions (Fig. 4C). To assess the contribution of elevated FTH1 levels to ferroptosis resistance during ECM-detachment, we engineered 786-O and MDA-MB-231 cells to be deficient in FTH1 (Fig. 4D). In both cell lines, shRNA targeting FTH1 was sufficient to sensitize ECM-detached cells to RSL3-mediated cell death (Fig. 4E). Similarly, shRNA targeting FTH1 sensitized ECM-detached cells to erastin (Fig. S4A) and cystine starvation (Fig. S4B) mediated cell death. Furthermore, the shRNA-mediated FTH1 deficiency caused a significant elevation in the levels of lipid ROS in ECM-detached cells exposed to RSL3 (Fig. 4F), erastin (Fig. S4C), or cystine starvation (Fig. S4D). Taken together, these findings suggest that elimination of ECM-detachment-mediated upregulation in FTH1 can sensitize ECM-detached cells to death by ferroptosis.

**Figure 4.**
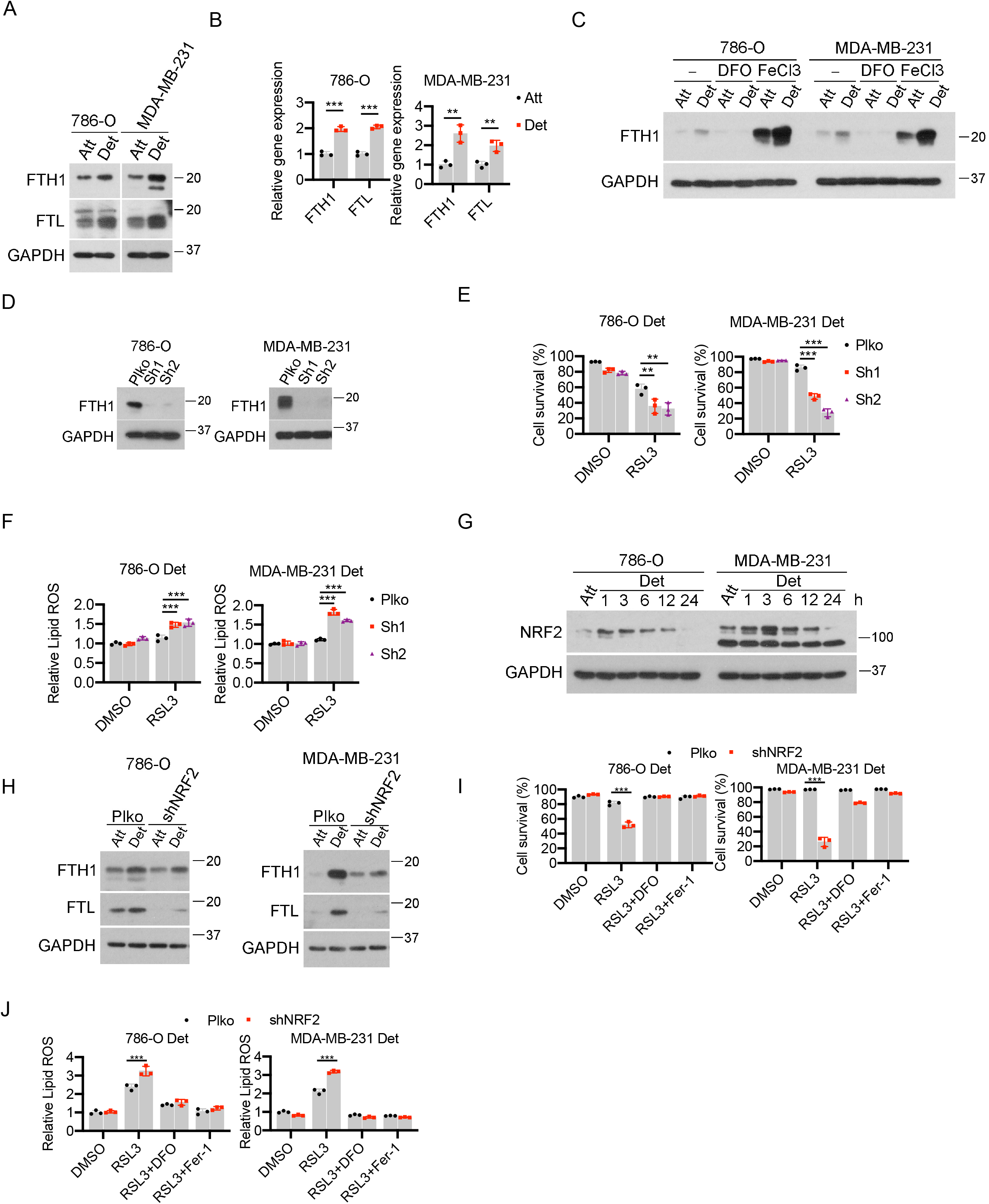
NRF2-FTH1 signaling contributes to ferroptosis resistance in ECM-detached cells. **(A)** Western blot detection of FTH1 and FTL protein expressions in cells detached for 24 h. **(B)** qRT-PCR detection of FTH1 and FTL gene expressions in cells detached for 24 h. **(C)** Western blot analysis of FTH1 in cells treated with 100 μM DFO or 100 μM FeCl_3_ for 24 h. **(D)** Western blot detection of FTH1 in cells stably expressing empty vector control (Plko) or FTH1 shRNAs. **(E)** Cell survival of detached cells treated by RSL3 (0.2 μM for 786-O, 1 μM for MDA-MB-231) for 24 h. **(F)** Lipid ROS of detached cells treated by RSL3 (0.2 μM for 786-O, 1 μM for MDA-MB-231) for 3 h. **(G)** Western blot detection of NRF2 expression in cells detached for different time points. **(H)** Western blot detection of FTH1 and FTL protein expressions in 786-O and MDA-MB-231 cells expressing empty control (Plko) or NRF2 shRNA (sh NRF2). **(I)** Cell survival of detached cells treated with RSL3 (0.2 μM for 786-O, 1 μM for MDA-MB-231) for 24 h. **(J)** Lipid ROS detection in detached cells treated with RSL3 (0.2 μM for 786-O, 1 μM for MDA-MB-231) for 3 h. For ferroptosis inhibition, 100 μM DFO or 2.5 μM Fer-1 was used. **p < 0.01, ***p < 0.001, two-tail student’s t test in B. **p < 0.01, ***p < 0.001, two-way ANOVA followed by Tukey test in E, F, I, J. Data are mean ±SD. Graphs represent data collected from a minimum of three biological replicates and all western blotting experiments were independently repeated a minimum of three times with similar results.

We next sought to determine the upstream regulators of FTH1 during ECM-detachment. Both FTH1 and FTL are known targets of the NRF2 transcription factor (Dodson et al., 2019; Kerins and Ooi, 2018), which is typically activated in order to neutralize excess oxidative stress (Rojo de la Vega et al., 2018; Sporn and Liby, 2012). Given that ECM-detached cells are often exposed to elevated oxidative stress (Davison et al., 2013; Hawk et al., 2018; Schafer et al., 2009), we investigated the levels of NRF2 when cells are grown in ECM-detached conditions. Indeed, we observed rapid, robust NRF2 activation in 786-O or MDA-MB-231 cells when grown in ECM-detachment (Fig. 4G). We next generated cells that were deficient in NRF2 (Fig. S4E) and found a substantial reduction in the abundance of both FTH1 and FTL protein (Fig. 4H) and mRNA (Fig. S4F) as a consequence of NRF2 shRNA. Moreover, when we assessed the sensitivity of NRF2-deficient cells to ferroptosis induction, we found that shRNA targeting NRF2 was sufficient to sensitize ECM-detached cells to RSL3 (Fig. 4I), erastin (Fig. S4G), or cystine starvation (Fig. S4H)-mediated cell death. Notably, cell death in each of these cases was blocked by treatment with DFO or Fer-1. Furthermore, NRF2-deficiency promoted lipid ROS accumulation in ECM-detached cells treated with RSL3 (Fig. 4J), erastin (Fig. S4I), or cystine starvation (Fig. S4J) which could also be blocked by either DFO or Fer-1. As such, these data suggest that antagonizing NRF2-mediated changes in FTH1 can sensitize ECM-detached cells to death by ferroptosis.

## Discussion

Our studies reveal that growth in ECM-detachment can substantially alter the sensitivity of cells to ferroptosis induction. ECM-detached cells are starkly resistant to ferroptosis triggered by distinct pharmacological stimuli (e.g. erastin, RSL3) and genetic changes (e.g. shRNA of GPX4). While ECM-detachment does promote enhanced cellcell contacts, diminished Yap-mediated expression of ACSL4, and reduced levels of PUFAs; these changes do not underlie the resistance of ECM-detached cells to ferroptosis induction. Instead, we found that ECM-detachment can cause profound changes in iron metabolism and demonstrated that the addition of excess, redox-active iron is sufficient to confer sensitivity to ferroptosis induction during ECM-detachment. We revealed that ECM-detached cells have both significant downregulation of TfR1-mediated iron uptake and upregulation of ferritin-related iron storage. Relatedly, we found that shRNA-mediated reduction of FTH1 was sufficient to sensitize ECM-detached cells to ferroptosis induction.

Our data suggest that as a consequence of ECM-detachment, cells undergo profound changes in iron metabolism that bestow significant resistance to ferroptosis induction. Notably, these changes in iron metabolism mirror other, previously defined changes in both glucose and glutamine metabolism and suggest that there are perhaps additional nutrients whose metabolism is altered during ECM-detachment (Grassian et al., 2011; Schafer et al., 2009). Furthermore, our data provide important nuance to understanding ferroptosis regulation. Previous studies have reported evidence that ECM-detachment can function as a trigger for the activation of ferroptosis due to deficient α6β4-mediated signal transduction (Brown et al., 2017). Thus, ECM-detachment could function to initiate ferroptosis in some cell types but ultimately those cells that survive acquire profound resistance to ferroptosis induction by other stimuli. In addition, the process of adaptation undertaken by cells that survive ECM-detachment may create *de novo* mechanisms of resistance or vulnerability.

Lastly, our data raise important questions about the capacity to utilize ferroptosis activation as a strategy against invasive or metastatic cancers. As discussed previously, there is broad interest in devising novel compounds that may target cancer cells for ferroptotic death (Gao et al., 2021; Hassannia et al., 2019; Koppula et al., 2022; Tarangelo et al., 2018; Wiernicki et al., 2022; Wu et al., 2022; Yi et al., 2020; Zhang et al., 2021). However, our data raise the distinct possibility that such approaches may need to account for the fact that ECM-detached cells, as a consequence of reprogramming of iron metabolism, may be resistant to efforts to cause ferroptosis. As such, therapeutic strategies that rely on ferroptosis induction may be hindered by the difficulty in killing ECM-detached cells. On the other hand, the reprogramming of iron metabolism during ECM-detachment may be a vulnerability that could be targeted to specifically eliminate cells upon loss of ECM-attachment. Future studies aimed at sensitizing cancer cells in ECM-detached conditions (e.g. circulating tumor cells) to ferroptosis may involve simultaneous treatment with ferroptosis activating compounds and agents that cause elevation in redox-active iron.

## Acknowledgements

We thank Veronica Schafer and all current/past Schafer lab members for helpful comments and/or valuable discussion. We thank the Flow Cytometry Facility and the Notre Dame Integrated Imaging Facility for experimental assistance. We thank Dr. Celeste Simon (UPenn) for the RCC-10 cell line. ZTS is supported by the National Institutes of Health/National Cancer Institute (R01CA262439), the Coleman Foundation, the Boler-Parseghian Center for Rare & Neglected Diseases at Notre Dame, the Malanga Family Excellence Fund for Cancer Research at Notre Dame, the College of Science at Notre Dame, the Department of Biological Sciences at Notre Dame, and funds from Mr. Nick L. Petroni. RGJ is supported by the National Institutes of Health/National Institute of Allergy and Infectious Diseases (R01AI165722), the Paul G. Allen Frontiers Group Distinguished Investigator Program, and the Van Andel Institute (VAI). We are also grateful for support and inspiration for this work from Mr. Christopher Kiergan and family.

## Author Contributions

JH, AMA, BPM, and RSM conducted experiments, analyzed data, and interpreted results. RS and RGJ assisted with the lipidomics and LC/MS experiments. JH and ZTS wrote the manuscript with feedback from all other authors. ZTS was responsible for conception/design of the project and overall study supervision.

## STAR Methods

### Resource Availability

#### Lead Contact

Further information and requests for resources and reagents should be directed to and will be fulfilled by the Lead Contact, Zachary T. Schafer.

#### Materials Availability

Plasmids and/or cell lines generated in this study are available upon reasonable request. Please contact the Lead Contact.

#### Data and Code Availability

The study did not generate/analyze any new datasets or code.

### Experimental Model and Subject Details

#### Cell lines

MDA-MB-231, 786-O, SUM159 and HeLa cells were from ATCC. RCC10 cell line was kindly provided by Dr. Celeste Simon (University of Pennsylvania Perelman School of Medicine). SUM159 cell line was maintained in Ham’s F12 (Gibco). All other cell lines were maintained in DMEM (Gibco). Culture medium was supplemented with 10% FBS (Gibco), 1% penicillin/streptomycin. Mycoplasma testing was routinely performed using mycoplasma PCR detection kit (LiliF, 102407-870) in all cell lines.

#### Reagents

Erastin (329600), RSL3 (SML2234), Ferrostatin-1 (SML0583) and Deferoxamine (DFO; D9533) were from Sigma. Methylcellulose (80080) was from Fisher Scientific.

#### Plasmids and retrovirus packaging

pQCXIH-Flag-YAP-S127A was from Addgene (#33092). pBabe-puro-TfR1-HA plasmid was generated by amplifying TfR1-HA from pcDNA3.2/DEST/hTfR-HA (Addgene, #69610) and inserting into pBabe-puro vector. For retrovirus packaging, 1.5 μg empty vector (EV) or plasmids carrying overexpressed genes were transfected to HEK293T cells together with 1.5 μg pCLAmpho. Transfection was performed using Lipofectamine 2000 (Invitrogen). Virus supernatant was collected 48 h after transfection and passed through 0.45 μm filter (EMD Millipore). Target cells were infected with virus supernatant in the presence of 10 μg/ml polybrene. Stable cell lines were selected using 750 μg/ml hygromycin (Sigma) for pQCXIH or 2 μg/ml puromycin (Invitrogen) for pBabe-puro.

#### Lentivirus mediated shRNA

For inducible GPX4 knockdown, shRNA sequence targeting human GPX4 (5’-GTGGATGAAGATCCAACCCAA-3’, TRCN0000046251) was cloned to Tet-Plko-puro according to the manual. Tet-pLKO-puro was a gift from Dmitri Wiederschain (Addgene, #21915). MISSION shRNA constructs for human FTH1 (NM_002032; clone nos TRCN0000029432, TRCN0000029433), human NRF2 (NM_006164; clone no TRCN0000273494), in the puromycin-resistant pLKO vectors were purchased from Sigma-Aldrich.

For packaging of lentivirus, HEK293T cells were transfected with 2 μg target DNA along with the packaging vectors psPAX2 (2.5 μg) and pCMV-VSV-G (0.5 g) using Lipofectamine 2000 (Invitrogen). Virus supernatant was collected 48 h after transfection and filtered through a 0.45 μm filter (EMD Millipore). Cells were infected with virus supernatant in the presence of 10 μg/ml polybrene. Stable populations of cells were selected using 2 μg/ml puromycin (Invitrogen).

### Method Details

#### Western blot

After treatment, both attached and detached cells are washed once with 2 ml PBS. ECM-attached and detached cells were lysed in RIPA buffer (20 mM Tris-HCl (pH 7.5), 150 mM NaCl, 1 mM Na2EDTA, 1 mM EGTA, 1% NP-40, 1% sodium deoxycholate) supplemented with 1 μg/ml aprotinin, 5 μg/ml leupeptin, 20 μg/ml phenylmethylsulfonyl fluoride (PMSF) and HALT phosphatase inhibitor mixture (Thermo Scientific). For attached cells, cells are lysed directly on 6-well plate. For detached cells, cells are resuspended in RIPA buffer. Cell lysates are sonicated with 1-s pulse on and 1-s pulse off for 8 s using Branson sonifier at 10% amplitude. After sonication, cell lysates were cleared by centrifuging at 13,000 rpm for 15 min. Supernatants were quantified by BCA assay (Pierce Biotechnology, Waltham, MA, USA) and normalized to same protein concentrations. 4X Lamelli sample buffer was added and heated at 95°C for 10 min. After SDS-PAGE, transfer and blocking, membranes were probed with primary antibodies at 4°C overnight. Horseradish peroxidase (HRP) linked secondary antibody incubation followed by chemiluminescent detection using SignalFire ECL Reagent (CST; 6883) was used to detect the blots. The following antibodies were used for western blotting: GPX4 (Abcam; ab41787; 1:2000), GCLC (Abcam; ab190685; 1:5000), ACSL4 (Santa Cruz biotechnology; sc-271800; 1:1000), GAPDH (CST; 5174S; 1:5000), FTH1 (CST; 3998S; 1:1000), FTL (Proteintech; 10727-1-AP; 1:1000), FSP1 (Proteintech; 20886-1-AP; 1:1000), xCT (CST; 12691), p-Yap S127 (CST; 13008T; 1:1000), HA-Tag (Biolegend; 901501; 1:2000), Flag-Tag (CST; 2368S; 1:1000), TfR1 (CST; 13113S; 1:1000).

#### Poly-HEMA coating

Poly-HEMA (Sigma; P3932) stock solution was made by dissolving 6 g Poly-HEMA in 1 L 95% ethanol (stir overnight at room temperature). For 6 well plates, add 1 mL poly-HEMA stock solution to each well, for 12 well plates add 0.5 mL poly-HEMA stock solution to each well. Keep plates in 37°C incubator for about one week until all liquid evaporates. Store coated plates at room temperature.

#### PI staining for cell viability

Cells plated on 12 well plates (8×10^4^ cells per well) under attached or detached (Poly-HEMA) conditions were treated with different compounds for indicated time. At the end of treatment, for attached cells, to include any floating dead cells, culture medium was collected. Cells were then washed with 300 μl PBS and PBS was combined together with culture medium. Cells were trypsinized and trypsin neutralized with the collected culture medium. For detached cells, cells were spun down and washed with 300 μl PBS. After trypsinization, trypsin was neutralized with complete medium. Both trypsinized attached and detached cells were spun down and resuspended in 180 μl PBS containing 2 μg/ml prodidium iodide (PI) (Sigma; P4170) and detected by flow cytometry (BD fortessa X-20). Single cells were gated and cell survival was measured by analyzing PI-negative and FSC-A high cells.

#### C11-Bodipy 581/591 staining

After treatment, both attached and detached cells were trypsinized to single cells as described in PI staining. Cells were then stained in 300 μl 1 μM C11-Bodipy 581/591 (Invitrogen; D3861) dissolved in PBS at 37 °C for 30 min. Cells were spun down, resuspended in 180 μl PBS and detected by flow cytometry. Live single cells were gated and geometry mean of FITC intensity was quantified. Relative lipid ROS was then calculated.

#### Calcein staining for labile iron

After treatment, both attached and detached cells were trypsinized to single cells. Cells were resuspended in 500 μl PBS containing 100 nM Calcein-AM (Thermo Fisher; C3099) and kept at 37 °C for 15 min. Cells were washed once with PBS and separated into two tube. Tube 1 was resuspended in 200 μl PBS and tube 2 was resuspended in 200 μl PBS with 100 μM deferiprone (Sigma; 379409). Both tubes were kept at 37°C for 30 min. Calcein fluorescence was detected by flow cytometry and difference in fluorescent intensity between tube 2 and tube 1 was used to indicate labile iron pool.

#### Transferrin uptake assay

After treatment, both attached and detached cells were balanced for 1 h in 500 μl serum-free DMEM containing 25 mM HEPES, pH7.4 with 1% BSA. Alexa fluor 488 conjugated human transferrin (Invitrogen; T13342) (5 mg/ml stock solution in PBS) was added at 20 μg/mL, kept at 37 for 30 min. Cells were washed with PBS and trypsinized to single cells. To remove surface-bound transferrin, cells were washed twice with pre-chilled PBS (pH 5.0). Transferrin uptake was determined by flow cytometry.

#### Real time PCR

Cells were plated on attached or detached conditions for 24 h. For mRNA extraction, RNeasy mini kit (Qiagen; 74104) was used. For attached cells, cells were lysed directly on plates. For detached cells, cells were spun down and lysed. Total mRNA was extracted according to manufacturer’s instructions. Reverse transcription was performed using reverse transcription supermix (Bio-rad; 1708840) and resulted cDNAs were further utilized for RT-PCR using SYBR green supermix (Biorad; 1725271). The following primers were used:

Human GPX4, Forward: 5’-ggagccagggagtaacgaag-3’; Reverse: 5’-cacttgatggcatttcccagg-3’

Human FTL, Forward: 5’-cagcctggtcaatttgtacct-3’; Reverse: 5’-gccaattcgcggaagaagtg-3’ Human FTH1, Forward: 5’-ccagaactaccaccaggactc-3’; Reverse: 5’-gaagattcggccacctcgtt-3’

Human TFRC, Forward: 5’-ggctacttgggctattgtaaagg-3’; Reverse: 5’-cagtttctccgacaactttctct-3’

Human ACSL4, Forward: 5’-actggccgacctaagggag-3’; Reverse: 5’-gccaaaggcaagtagccaata-3’

Human 18S, Forward: 5’-ggcgccccctcgatgctcttag-3’; Reverse: 5’-gctcgggcctgctttgaacactct-3’

#### Lipidomics and mass spectrometry

Lipids were extracted from cells by Bligh Dyer extraction. For adherent cells, plates were first scraped into ice cold methanol, then transferred to a tube containing chloroform. Suspension cells were lysed directed in chloroform:methanol (1:1, v/v). To induce phase separation, water was added to achieve a final chloroform:methanol:water ratio of 2:2:1.8 (v/v). The lipid containing organic layer was collected, dried in a speedvac, and resuspended in 1:1 (v/v) isopropanol:acetonitrile. 100uL of each sample was used to make a pooled sample for lipid identification. Lipidomics analysis was collected by LC/MS using an ID-X Orbitrap mass spectrometer (Thermo Scientific)> 2uL of reconstituted extract was injected for liquid chromatography separation using a C30 column (27826-152130, Thermo) maintained at 50°C. Lipids were eluted with a gradient (A: 60% acetonitrile, B: 90% isopropanol, 8% acetonitrile, both with 10mM ammonium formate and 0.1% formic acid) of 0-30min starting at 75%A, 1-3min ramp from 75%A to 60%A, 3-19min from 60%A to 25%A, 19-20.5min from 25%A to 10%A, 20.5-28min from 10%A to 5%A, 28-28.1min from 5%A to 0%A, and ending with an isocratic hold at 0%A from 28.1-30min at 400uL/min. Lipids were detected in ESI+ positive mode (spray voltage of 3250 V, sheath gas: 40 a.u., aux gas: 10 a.u., sweep gas: 1 a.u., ion transfer tube: 300°C, vaporizer: 275°C). Experimental replicates were analyzed in MS1 only at a resolution of 240,000 FWHM. Pooled samples were used for lipid ID via data dependent MS3 method that triggers all ions reaching an intensity threshold 10x above a blank for HCD fragmentation. An HCD MS2 fragment of 184, a diagnostic ion for phosphatidyl choline, triggered an additional MS2 fragmentation scan using CID to gain information on acyl chain composition. A neutral loss of acyl chains triggered MS3 scan in HCD to gain further structural information. ddMS3 samples. ddMS3 data from pooled samples was used for lipid annotation in LipidSearch (Thermo, v5.0). Annotated lipids were then used to generate a mass list (accurate mass at retention time) for Compound Discoverer (Thermo, v3.3), which was used for peak picking and integration of experimental replicates. Differential abundance of lipids was accomplished with Metabolanlyst (v5.0) with an FDR set at 0.05.

### Quantification and Statistical Analysis

Data were presented as mean ±SD. All graphs and statistical analysis were made in Graphpad Prism (8.0.0). Student two-tailed unpaired *t*-test was used for two-group comparison and two-way analysis of variance (ANOVA) followed by Tukey test was used for multi-group comparisons. Details for each figure are included in the Figure Legends.

---

**Figure S1.**
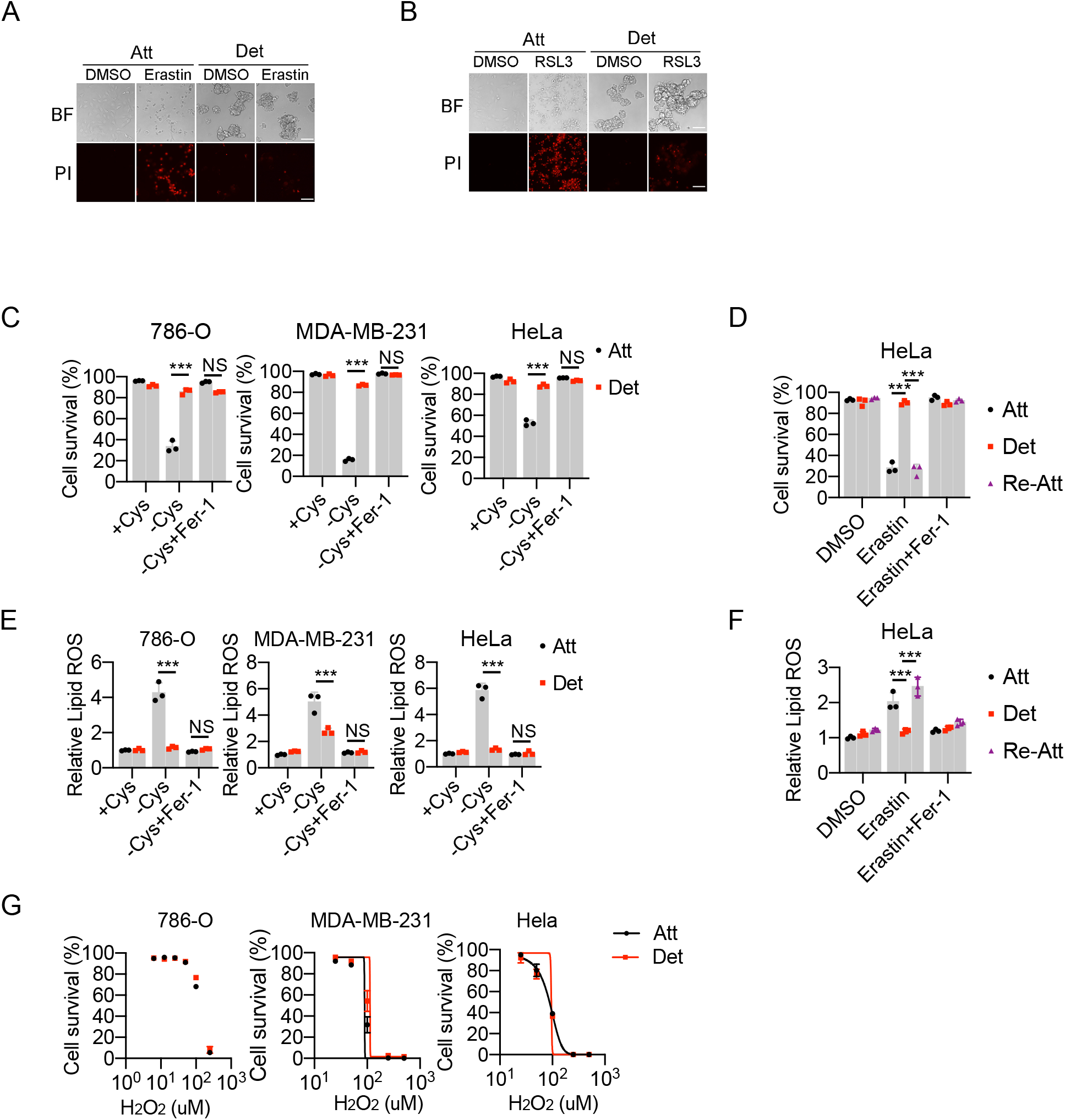
ECM-detached cells are resistant to ferroptosis induction. Related to Figure 1. (A-B) Representative images of 780-O cells plated on Att or Det conditions treated with erastin (5 μM, 24 h) (A) or RSL3 (0.2 μM, 16 h) (B). BF, bright field; PI, prodidium iodide. Scale bar, 50 μm. (C) Survival of cells under cystine starvation (-Cys) for 24 h. 2.5 μM Fer-1 was used to block ferroptosis. (D) Survival of Hela cells treated with 5 μM erastin under attached (Att), detached (Det) or re-attached (Re-Att) conditions for 24 h. For re-attachment, cells were first plated on poly-HEMA coated plate for 12 h and switch to attachment plates for erastin treatment. (E-F) Lipid ROS detection of cells with cystine starvation for 16 h (E) or 5 μM erastin treatment (12 h) (F). (G) Survival of cells treated with H2O2 for 24 h. ***p < 0.001, NS, not significant, two-way ANOVA followed by Tukey test. Data are mean +/− SD. Graphs represent data collected from a minimum of three biological replicates.

**Figure S2.**
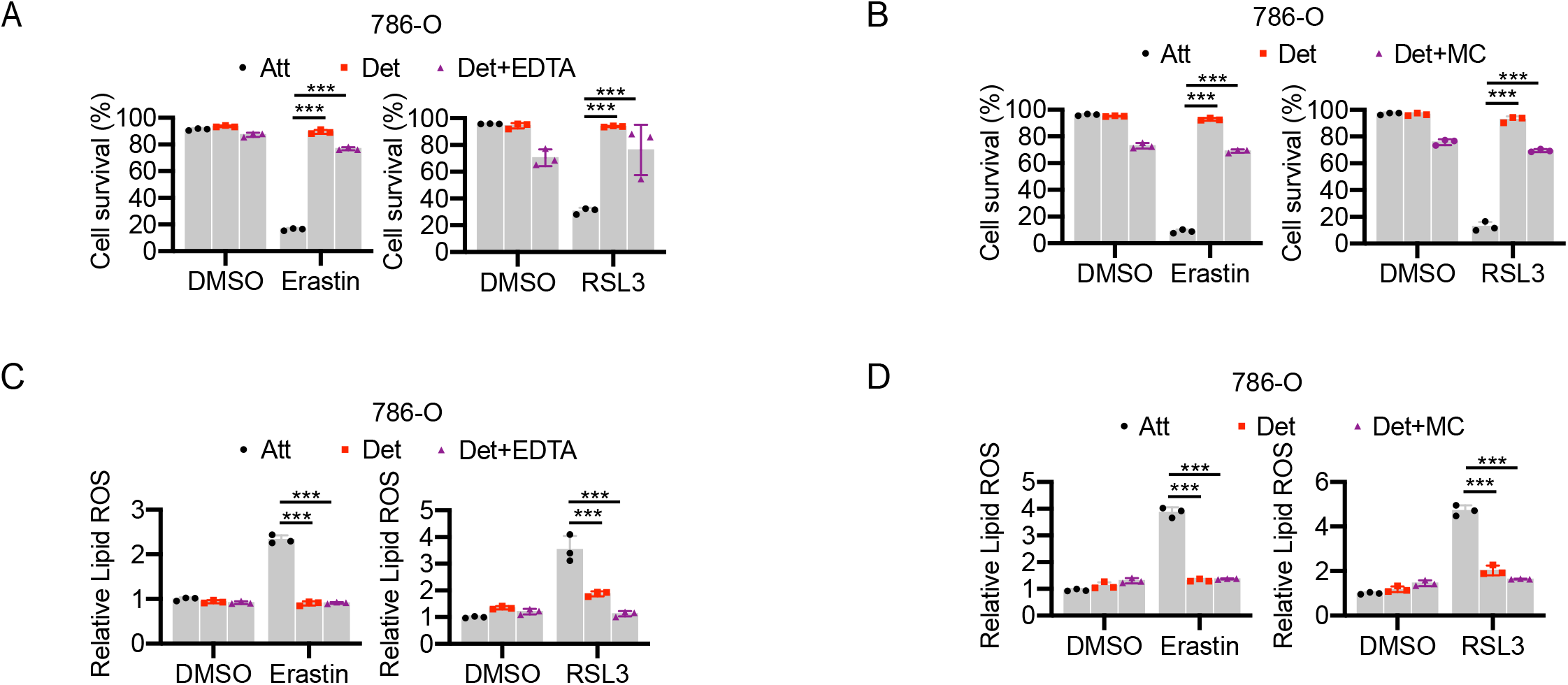
Disruption of cell cluster formation does not sensitize ECM-detached cells to ferroptosis induction. Related to Figure 2. (A-B) Survival of cells treated with 5 μM erastin (A) or 0.2 μM RSL3 (B) for 24 h. 2 mM EDTA or 0.5% Methylcellulose was added to block the formation of cell clusters. (C-D). Lipid ROS detection of cells treated with 5 μM erastin (12 h) (C) or 0.2 μM RSL3 (3 h) (D). 2 mM EDTA or 0.5% Methylcellulose was added to block the formation of cell clusters. ***p < 0.001, two-way ANOVA followed by Tukey test. Data are mean +/− SD. Graphs represent data collected from a minimum of three biological replicates.

**Figure S3.**
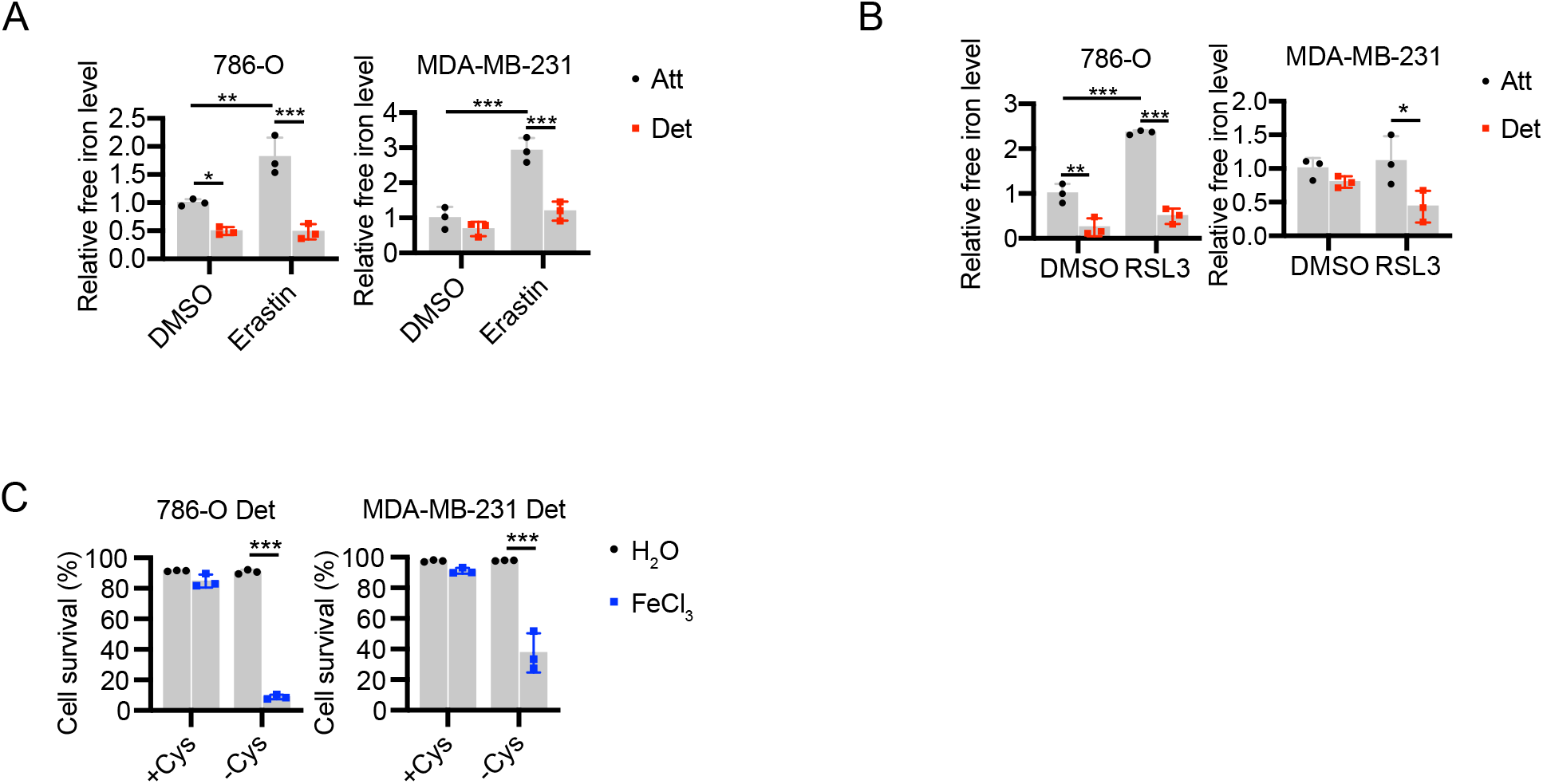
Iron uptake is reduced in ECM-detached cells. Related to Figure 3. (A) Free iron level detection in cells treated with either DMSO or 5 μM Erastin for 12 h. (B) Free iron level detection in cells treated with either DMSO or 0.2 μM RSL3 for 3 h. (C) Survival of detached cells with cystine starvation under either H2O or FeCl3 (100 μM) treatment for 24 h. *p < 0.05, **p < 0.01, ***p < 0.001, two-way ANOVA followed by Tukey. Data are mean +/− SD. Graphs represent data collected from a minimum of three biological replicates.

**Figure S4.**
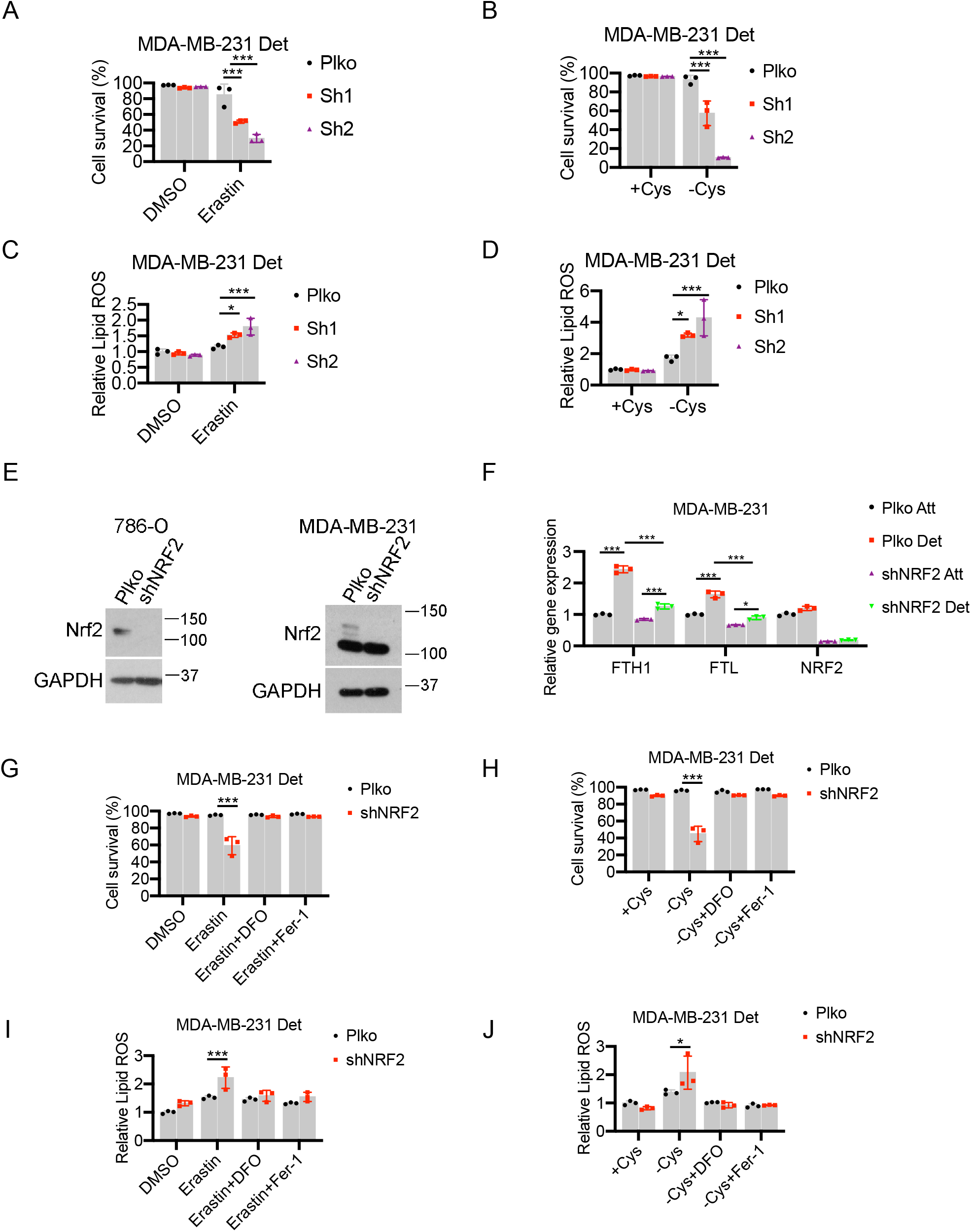
NRF2-FTH1 signaling contributes to ferroptosis resistance in ECM detached cells. Related to Figure 4. (A-B) Survival of detached MDA-MB-231 cells expressing either control (Plko) or shRNAs against FTH1 treated with 10 μM erastin (A) or cystine starvation for 24 h (B). (C-D) Lipid ROS detection of cells treated with 10 μM erastin (C) or cystine starvation (D) for 12 h. (E) Western blot analysis of NRF2 expression in 786-O and MDA-MB-231 cells expressing plko or shNRF2. (F) Quantitative real-time PCR analysis of FTH1, FTL and NRF2 expressions in MDA-MB-231 cells. (G-H) Cell survival of detached cells treated with 10 μM erastin (G) or cystine starvation (H) for 24 h. (I-J) Lipid ROS detection of cells treated with 10 μM erastin (I) or cystine starvation (J) for 12 h. For ferroptosis inhibition, 100 μM DFO or 2.5 μM Fer-1 was used. *p < 0.05, **p < 0.01, ***p < 0.001, two-way ANOVA followed by Tukey test. Data are mean +/− SD. Graphs represent data collected from a minimum of three biological replicates and all western blotting experiments were independently repeated a minimum of three times with similar results.

